# The males of the parasitoid wasp, *Nasonia vitripennis*, can identify which fly hosts contain females

**DOI:** 10.1101/2021.04.06.438549

**Authors:** Garima Prazapati, Ankit Yadav, Anoop Ambili, Abhilasha Sharma, Rhitoban Raychoudhury

## Abstract

The reproductive success of a male is limited by the number of females it can mate with. Thus, males deploy elaborate mate-finding strategies to maximize access to females. In the haplodiploid wasp genus, Nasonia, which are parasitoids of cyclorrhaphous flies, mate-finding is restricted to the natal patch, where males compete for access to females. This study investigates whether there are any additional mate finding strategies of males, especially, whether they can identify the presence of adult females which are still inside the fly host. Results reveal that only one out of the four species, N. vitripennis, can distinguish which hosts specifically have adult female wasps indicating a species-specific unique mate-finding capability. Behavioral assays revealed that the cues used by N. vitripennis males are olfactory in nature and not auditory or visual. GC-MS analyses show that these olfactory cues are female-specific cuticular hydrocarbons (CHCs), possibly emanating from within the fly puparium. Further assays indicated that N. vitripennis males can also detect differences in the concentrations of compounds to identify female-specific cues from male-specific ones. This study, therefore, uncovers a previously unknown mate-finding strategy in one of the most widely studied parasitoid wasp.

## Introduction

In most sexually reproducing organisms, male reproductive success is limited by the number of fertile females it can mate. In contrast, female reproductive success is mainly limited by the number of eggs produced (Bateman, 1948). This difference necessitates distinct reproductive strategies for both (Gross, 1996). For a male, the ideal reproductive strategy involves rapid sexual maturation and access to many fertile females (Kappeler, 2012; Muller and Thompson, 2012). To achieve this, males have evolved several mate-finding strategies (Andersson, 1994). In male parasitoid wasps, finding many females is relatively easy as most species show a female-biased sex ratio (Godfray, 1994). Despite this, male parasitoid wasps adopt various mate-finding strategies to maximize individual fitness. These include the use of trail sex pheromone deposited by females in *Aphelinus asychis* (Fauvergue *et al*., 1995), *Aphytis melinus* (Bernal and Luck, 2007), and *Trichogramma brassicae* (Pompanon *et al*., 1997). *Urolepis rufipes* use territorial markings (Cooper and King, 2015) and emergence sites of con-specific males (Wittman *et al*., 2016) to attract females. In some parasitoid wasps, mate-finding involves using chemical cues from the hosts themselves (Vinson, 1976). *Pimpla disparis* males use vibratory or acoustic cues emanating from developing wasps inside the gypsy moth (*Lymantria dispar*) host (Hrabar *et al*., 2012; Danci *et al*., 2014). *Cephalonomia tarsalis* (Collatz *et al*., 2009) use host-associated sex pheromones for finding mates whereas, *Lariophagus distinguendus* (Steiner *et al*., 2007) males use volatile cues, other than sex pheromones, to do so.

No specific mate-finding strategy has been uncovered in the pteromalid wasp *Nasonia*, one of the most extensively studied parasitic wasp (Mair and Ruther, 2019). The haplodiploid parasitoid wasp genus, *Nasonia*, comprises four species, *N. vitripennis, N. longicornis, N. giraulti*, and *N. oneida* (Raychoudhury *et al*., 2010), and parasitizes on cyclorrhaphous fly pupae. The female locates a suitable fly pupa (host), drills through the puparium by its ovipositor, paralyzes the fly pupa by injecting it with venom, and then lays eggs (Whiting, 1967). The entire holometabolous life-cycle, from eggs to adults, happens inside the host, and the adults emerge by chewing an emergence hole through the host’s puparium. Although all four *Nasonia* species have a female-biased offspring sex ratio, curiously, it is the male which usually emerges first (Cousin, 1933) and waits around for emerging females (Werren, 1980). Mating happens quickly, and the females then fly away in search of newer hosts to parasitize. Little is known whether the males possess any other strategy to find mates or are even capable of actively seeking out females beyond hanging around the emergence holes. However, several biological features of *Nasonia* indicate that males can be under relatively intense selection pressure to evolve strategies looking for females beyond their natal host. Nasonia females parasitize hosts available as a patchy resource, and the female often parasitizes as many as she can (Godfray, 1997). Therefore, most of the emerging progeny are relatively close to each other and within reach of any emerged male.

Moreover, the males are reproductively mature as soon as they emerge with a full complement of functional sperm (Chirault, 2016) and do not leave their natal patch (Van den Assem and Vernel, 1979). Since females mate only once (Grillenberger *et al*., 2008), males can fertilize many females. Males compete to access females by aggressively defending the host puparium from which they emerge and never leaving the natal host (Leonard and Boake, 2006). This intrasexual aggression can also be a trigger for additional strategies for finding mates. One such possibility is the ability to detect hosts about to release adult females. There is some evidence that males can recognize parasitized fly hosts as they spend significantly more time on them than unparasitized ones (King *et al*., 1969). However, what remains unknown is whether this ability extends to finding out whether a parasitized host will have females inside to mate with, as *Nasonia* is a haplodiploid wasp, and some hosts might have all-male broods. This study conducts a comprehensive investigation of this potential mate-finding strategy across the four *Nasonia* species by determining their preference for differentially parasitized fly hosts of various development stages. We also determine the nature of the cues (auditory, visual, or olfactory) utilized by the males and identify the olfactory cues’ chemical nature by GC-MS. We find a species-specific mate finding strategy that depends on males’ ability to detect different concentrations of chemicals involved in olfaction.

## Results

### *Nasonia* males can detect parasitized hosts

We first established whether *Nasonia* males can detect parasitized hosts within a given patch which also contains unparasitized ones. Males were given a choice between two-day old parasitized (HwL) and unparasitized hosts. As figure 1 (a) indicates, males of all four species can detect which hosts are parasitized as they spent significantly more time on them (*p* < 0.01, *r* = 0.8 for *N. vitripennis*; *p* < 0.01, *r* = 0.8 for *N. longicornis*; *p* < 0.01, *r* = 0.6 for *N. giraulti* and *p* < 0.01, *r* = 0.8 for *N. oneida*). However, each HwL has larval wasps inside it and is several days away from adult wasp eclosion which can extend well beyond the life span of an adult male. Hence, to test whether males can detect parasitized hosts which contain eclosed adults (HwAMF), a choice was given between such hosts and unparasitized ones. As figure 1 (b) shows, males of all the four species spent significantly more time on HwAMF (*p* < 0.001, *r* = 0.8 for *N. vitripennis*; *p* < 0.01, *r* = 0.6 for *N. longicornis*; *p* = 0.01, *r* = 0.4 for *N. giraulti* and *p* < 0.001, *r* = 0.8 for *N. oneida*). Thus, *Nasonia* males can identify hosts which contain larval as well as adult wasps. However, as mentioned above, detecting larval wasps will add little to the reproductive success of any male. Therefore, we gave a choice between these two types of hosts (HwL and HwAMF) to determine whether males have the capability to distinguish which host has adult wasps inside. Interestingly, *N. vitripennis* (*p* < 0.01, *r* = 0.5) and *N. oneida* (*p* = 0.01, *r* = 0.5) showed a preference for HwAMF (figure 2 a) but *N. longicornis* (*p* = 0.7, *r* = 0) and *N. giraulti* (*p* = 0.6, *r* = 0.1) did not. Thus, there is species-specific variation for this particular capability where *N. vitripennis* and *N. oneida* are able to identify adult wasps within the hosts but *N. longicornis* and *N. giraulti* cannot.

**Figure 1:**
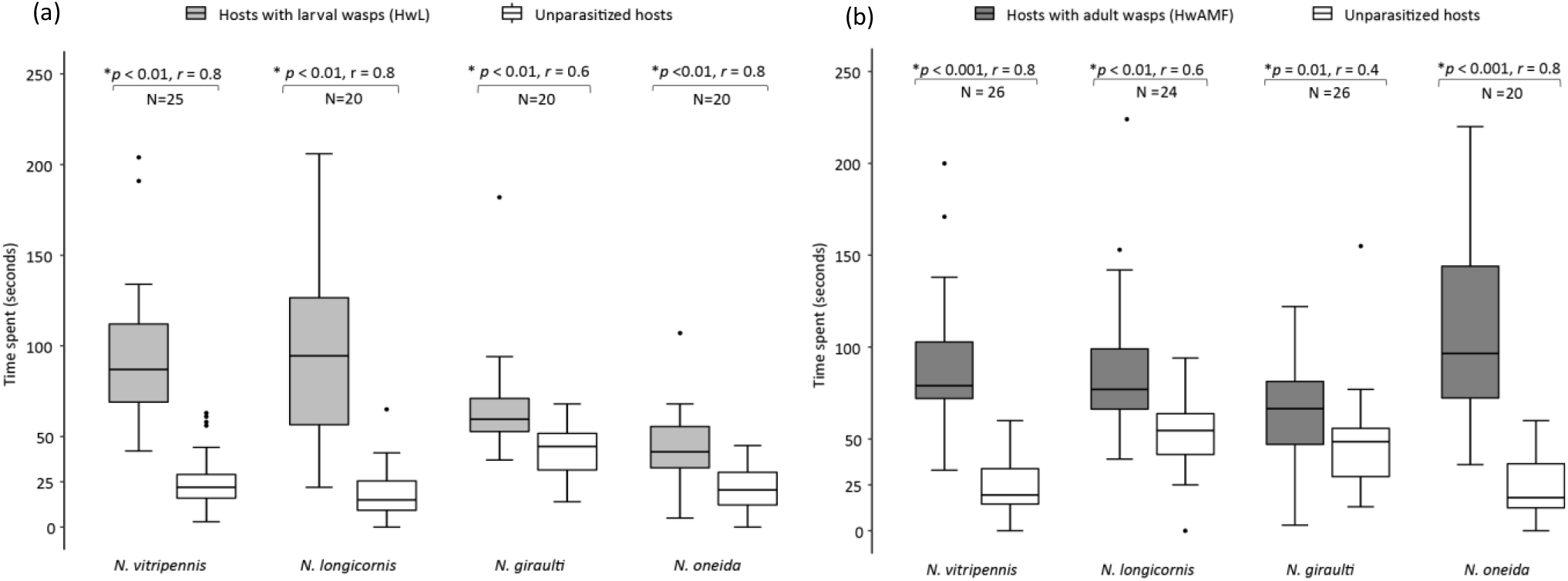
*Nasonia* males can detect parasitized hosts: **(a)** Average time spent by males of all the four species, on parasitized hosts containing larval wasps (HwL) versus unparasitized ones. Males of all the four species spend significantly more time on parasitized hosts indicating their ability to detect hosts with larval wasps inside. **(b)** Average time spent by males on parasitized hosts containing adult wasps (HwAMF) and unparasitized ones. Males of all the four species spend significantly more time on parasitized hosts indicating their preference for hosts with adult wasps inside. The numbers above the boxes represent the *p-*value and the sample size (N) for each species. In boxplots, the horizontal bold line represents the median, boxes represent 25% and 75% quartiles, whiskers denote 1.5 interquartile ranges and black dots depict outliers. Statistical significance levels shown are according to Wilcoxon signed-rank test (statistically significant at *p* < 0.05) with (*) denoting a significant *p*-value. Wilcoxon effect size (*r*) values range from *r* = 0.1 - < 0.3 (small effect), *r* = 0.3 - < 0.5 (moderate effect) and *r* >= 0.5 (large effect). Species names are given at the bottom.

**Figure 2:**
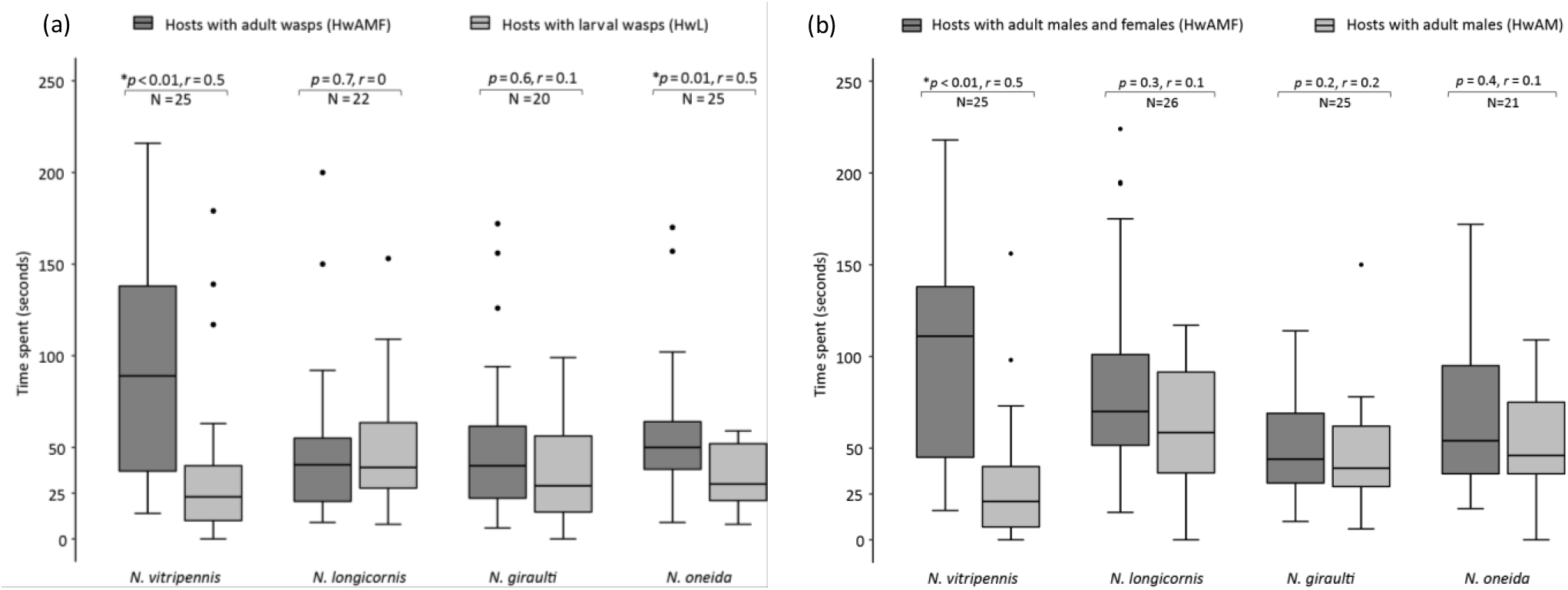
*N. vitripennis* males can detect adult females within hosts: **(a)** Average time spent by males of all the four species on parasitized hosts containing adult wasps (HwAMF) and those containing larval wasps (HwL). Males of *N. vitripennis* and *N. oneida* can distinguish between HwAMF over HwL, whereas, *N. longicornis* and *N. giraulti* cannot. **(b)** *N. vitripennis* males can distinguish between hosts with males and females (HwAMF) over those containing all - male broods (HwAM). *N. longicornis*, *N. giraulti* and *N. oneida* do not show this capability. Thus, *N. vitripennis* males can detect adult females still inside the hosts. The numbers above the boxes represent the *p-*value and the sample size (N) for each species. In boxplots, the horizontal bold line represents the median, boxes represent 25% and 75% quartiles, whiskers denote 1.5 interquartile ranges and black dots depict outliers. Statistical significance levels shown are according to Wilcoxon signed-rank test (statistically significant at *p* < 0.05) with (*) denoting a significant *p*-value. Wilcoxon effect size (*r*) values range from *r* = 0.1 - < 0.3 (small effect), *r* = 0.3 - < 0.5 (moderate effect) and *r* >= 0.5 (large effect). Species names are given at the bottom.

### *N. vitripennis* males can detect adult females within hosts

*Nasonia* being a haplo-diploid wasp can also reproduce via arrhenotokous parthenogenesis, where unfertilized eggs will give rise to males, resulting in all-male broods. Thus, the capability to detect hosts containing adult wasps will add to the fitness of a male only if it can detect which hosts will yield adult females. Thus, *Nasonia* males were given a choice between hosts which had all-male adult broods (HwAM) and those that had the adults of both sexes (HwAMF). Interestingly, as figure 2 b illustrates, only *N. vitripennis* showed a significant preference for the latter (*p* = 0.001, *r* = 0.6), while all the other three could not distinguish between them (*N. longicornis*, *p* = 0.3, *r* = 0.1; *N. giraulti*, *p* = 0.2, *r* = 0.2; *N. oneida*, *p* = 0.4, *r* = 0.1). Thus, *N. vitripennis* males are not only capable of detecting which host will yield adults, but they can also distinguish which ones will have females in them. This proficiency is not affected even by the presence of adult males inside. Next, we investigated the possible cues that males of *N. vitripennis* utilize to elicit this phenotype.

### *N. vitripennis* males do not use auditory and visual cues to detect adult females within hosts

One of the possible cues that males can use is the sound coming from inside the host as the adult wasps eclose before emerging from the host. To test this possibility, *N. vitripennis* males were given a choice between hosts with alive adult wasps inside, and other hosts with dead (freeze-killed) adult wasps inside, thereby removing auditory cue coming from adults. As shown in figure 3 a, males of *N. vitripennis* did not show a preference (*p* = 1, *r* = 0) for either type of host, indicating that they probably do not utilize auditory cues for detecting hosts with adult females inside.

**Figure 3:**
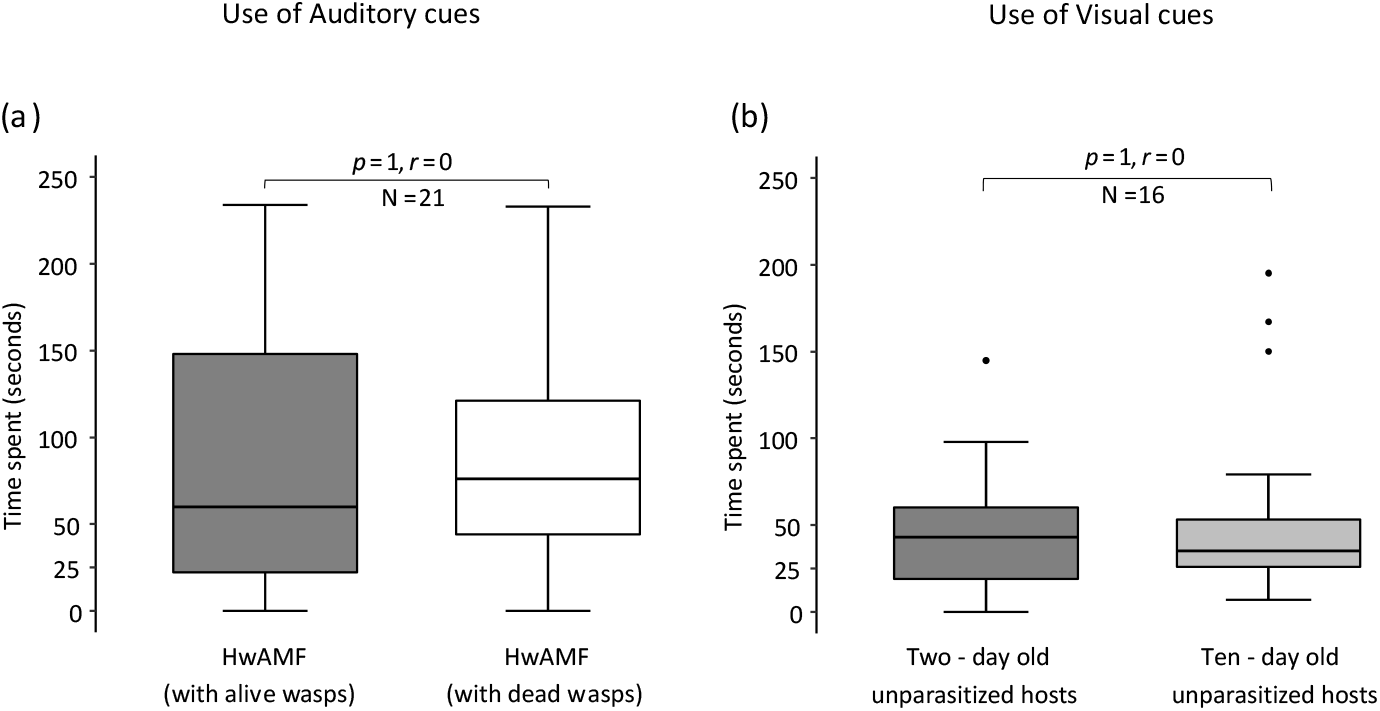
*N. vitripennis* do not utilize auditory and visual cues to detect adult females within hosts: **(a)** No significant difference was found between average time spent by males on HwAMF with alive wasps and those with dead wasps. Hence, males do not utilize auditory cues. **(b)** No significant difference was found between average time spent by males on the puparial halves of two - day old and ten - day old unparasitized hosts. Hence, males do not utilize visual cues. The numbers above the boxes represent the *p-*value and the sample size (N), respectively. In boxplots, the horizontal bold line represents the median, boxes represent 25% and 75% quartiles and whiskers denote 1.5 interquartile ranges and black dots depict outliers. Statistical significance levels shown are according to Wilcoxon signed-rank test (statistically significant at *p* < 0.05) with (*) denoting a significant *p*-value. Wilcoxon effect size (*r*) values range from *r* = 0.1 - < 0.3 (small effect), *r* = 0.3 - < 0.5 (moderate effect) and *r* >= 0.5 (large effect).

Another possible cue that males can utilize is the physiochemical changes happening in the host as it develops. The puparium, which is the host pupa’s outer casing, undergoes perceptible darkening with time (Sinha and Mahato, 2016) and can be a visual cue for discrimination. We investigated whether such puparial darkening can act as a visual cue by giving them a choice between puparial halves obtained from the anterior end of two-day-old unparasitized hosts and those obtained from ten-day-old unparasitized hosts (a day before adult fly eclosion, hence, maximally darkened. As figure 3 (b) shows, males did not distinguish between these two types of puparial halves and spent nearly equal time on both (*p* = 1, *r* = 0). Therefore, puparial darkening is not a cue utilized by *N. vitripennis* males.

### *N. vitripennis* males use olfactory cues to detect adult females within hosts

*Nasonia* uses several olfactory cues during courtship (Van den Assem *et al*., 1980), mate-choice (Ruther *et al*., 2009; Ruther *et al*., 2011), and even for species-recognition (Mair *et al*., 2017; Buellesbach *et al*., 2013; Buellesbach *et al*., 2018). These olfactory cues can include cuticular lipids acting as contact sex-pheromones and other as yet unknown semiochemicals (Mair and Ruther, 2019). The ability of *N. vitripennis* females to recognize and assess the quality of a parasitized host is hypothesized to involve chemosensory cues (Blaul *et al*., 2014; King and Rafai, 1970). However, whether *N. vitripennis* males use similar cues is not known.

There are two possible sources of olfactory cues that a male can utilize to locate adult females within hosts. The first is any olfactory cues left behind by a female during parasitization while the second can be any olfactory cues emanating from the wasps within the host. To test the first possibility, an unparasitized host was partially embedded in a foam plug, with only the anterior half exposed to the female for parasitization for 48 hours (SI - Figure S3). Males were given a choice between exposed puparial halves of such parasitized hosts (HwL) and those that were not exposed to females. Using just the puparial halves of the same age ensured that no other cues (visual, auditory, *etc.*) would influence the choice. As figure 4 (a) shows, males spent significantly more time (*p* < 0.001, *r* = 0.8) on the puparial halves exposed to females, indicating that they can perceive any olfactory cues left behind by the female wasp.

**Figure 4:**
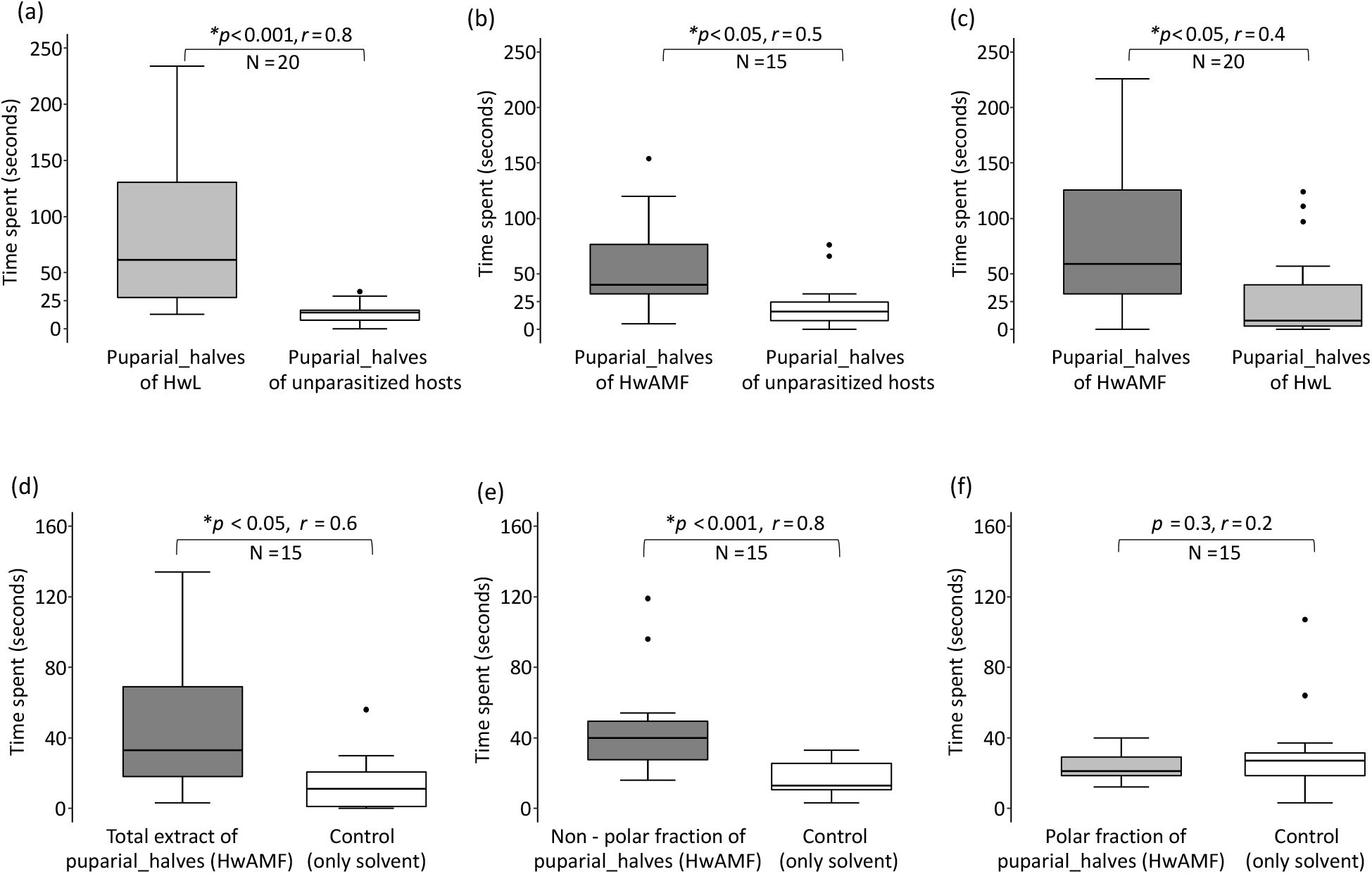
*N. vitripennis* males utilize olfactory cues to detect adult females within hosts: **(a)** Males prefer parasitized puparial halves of HwL (hosts containing larval wasps) over those of hosts containing fly pupa. **(b)** Males prefer puparial halves of HwAMF (hosts containing adult wasps) over unparasitized hosts. **(c)** Average time spent by the males on puparial halves of HwAMF versus HwL. Males spend significantly more time on the former. **(d)** Average time spent by males on the total extract of puparial halves of HwAMF versus the control poured with the solvent (DCM). Males prefer the extract, indicating that they utilize olfactory cues. **(e)** Average time spent by males on the non-polar fraction of the extract (enriched for CHCs) versus control. Males show significant preference for the non-polar fraction. **(f)** Average time spent by males on the polar fraction of the extract versus control. Males show no preference towards either. Thus, the nature of olfactory cues utilized by the *N. vitripennis* males is non-polar, usually enriched for CHCs The numbers above the boxes represent the *p-*value and the sample size (N), respectively. In boxplots, the horizontal bold line represents the median, boxes represent 25% and 75% quartiles and whiskers denote 1.5 interquartile ranges and black dots depict outliers. Statistical significance levels shown are according to Wilcoxon signed-rank test (statistically significant at *p* < 0.05) with (*) denoting a significant p-value. Wilcoxon effect size (*r*) values range from *r* = 0.1 - < 0.3 (small effect), *r* = 0.3 - < 0.5 (moderate effect) and *r* >= 0.5 (large effect).

Next, we tested whether the eclosed wasps were emanating any olfactory cues. The puparium of a host is a porous structure (Yoder and Denlinger, 1991), and, hence, males can perceive any olfactory cues coming from within. However, the host fly ecloses within eleven days at 25 °C, while the wasps eclose by the fourteenth day. Therefore, the chronological age of the two types of hosts differs from the developing insects’ physiological age. To minimize the effect of this disparity, males were given a choice between the puparial halves obtained from the anterior half of the two types of hosts *i.e*., HwAMF and those containing adult fly (ten-day old). This ensured that the males were choosing between two types of puparial halves that had the maximum physiological age. Males spent significantly more time (*p* < 0.05, *r* = 0.5; figure 4 b) on the puparial halves of HwAMF, indicating that either the olfactory cues deposited by the parasitizing female persist throughout the life-cycle of the wasps or additional olfactory cues are emanating from the adult wasps within. Interestingly, when males were given a choice between the puparial halves of hosts containing adult wasps (HwAMF) and those containing larval ones (HwL), males prefer the former (*p* < 0.05, *r* = 0.4; figure 4 c). This indicates that either males can perceive any additional olfactory cues emanating from the adult wasps inside or these adults are probably producing the cues at a higher concentration. It seems logical that males can be under selection to detect the latter source, as perceiving hosts containing adult wasps would substantially increase the chance of encountering mates.

To confirm whether the cues utilized by *N. vitripennis* males is olfactory in nature, the puparial halves of HwAMF were dipped in dichloromethane (DCM) for 20 minutes and the obtained extract was pipetted out in a separate glass vial. Male preference was tested towards the extract poured over puparial halves of unparasitized hosts against those poured over with only DCM. As figure 4 (d) indicates, males spent significantly more time on the puparial halves with the extract (*p* < 0.05, *r* = 0.6). This confirms that they utilize olfactory cues to identify hosts containing potential mates, since all other cues (visual and auditory) that could otherwise influence such a choice were absent.

To identify the chemical nature of the olfactory cues, the polar and non-polar fractions were separately enriched through column chromatography (see Methods).The polar fraction of the extract usually contains sex-pheromones and polar lipids such as cholesterol, free fatty acids, *etc*. The non-polar component contains lipids such as cuticular-hydrocarbons (CHCs) (Mair *et al*., 2017; Carlson *et al*., 1998; Carlson *et al*., 1999). Male preference was tested towards each of these fractions and as figure 4 (e) shows, they prefer the non-polar fraction (*p* < 0.01, *r* = 0.8) and not the polar fraction (*p* = 0.3, *r* = 0.2; figure 4 f). This indicates that the source of the olfactory cues is present in the non-polar fraction and since it is enriched for CHCs, these could be the source of the olfactory cue.

### *N. vitripennis* males utilize cuticular hydrocarbons (CHCs) of females, as olfactory cues, to detect adult females within hosts

GC-MS of the non-polar fraction obtained from HwAMF and HwAM identified an array of long-chain saturated as well as unsaturated hydrocarbons with carbon chain lengths ranging from *n*C25- *n*C37, mostly consisting of *n*-alkanes, alkenes, mono-, di-, tri-, and tetra-methyl alkanes (Figure 5; SI - Table S1). The most abundant compound was Hentriacontane (*n*C-31) in both HwAMF and adult females, Nonacosane (*n*C-29) in HwAM and 7-methyltriacontane (MeC31 (7-)) in adult males. However, a comparative assessment of the CHC profiles of HwAMF and HwAM shows no detectable compositional change between the two as they share all the 53 compounds (SI - Table S1). A principal component analysis shows no clear separation between the two profiles (Figure 6) unlike the adult male and female CHC profiles which also have no detectable compositional change, as noted by previous studies (Buellesbach *et al*., 2013; Buellesbach *et al*., 2018).

**Figure 5:**
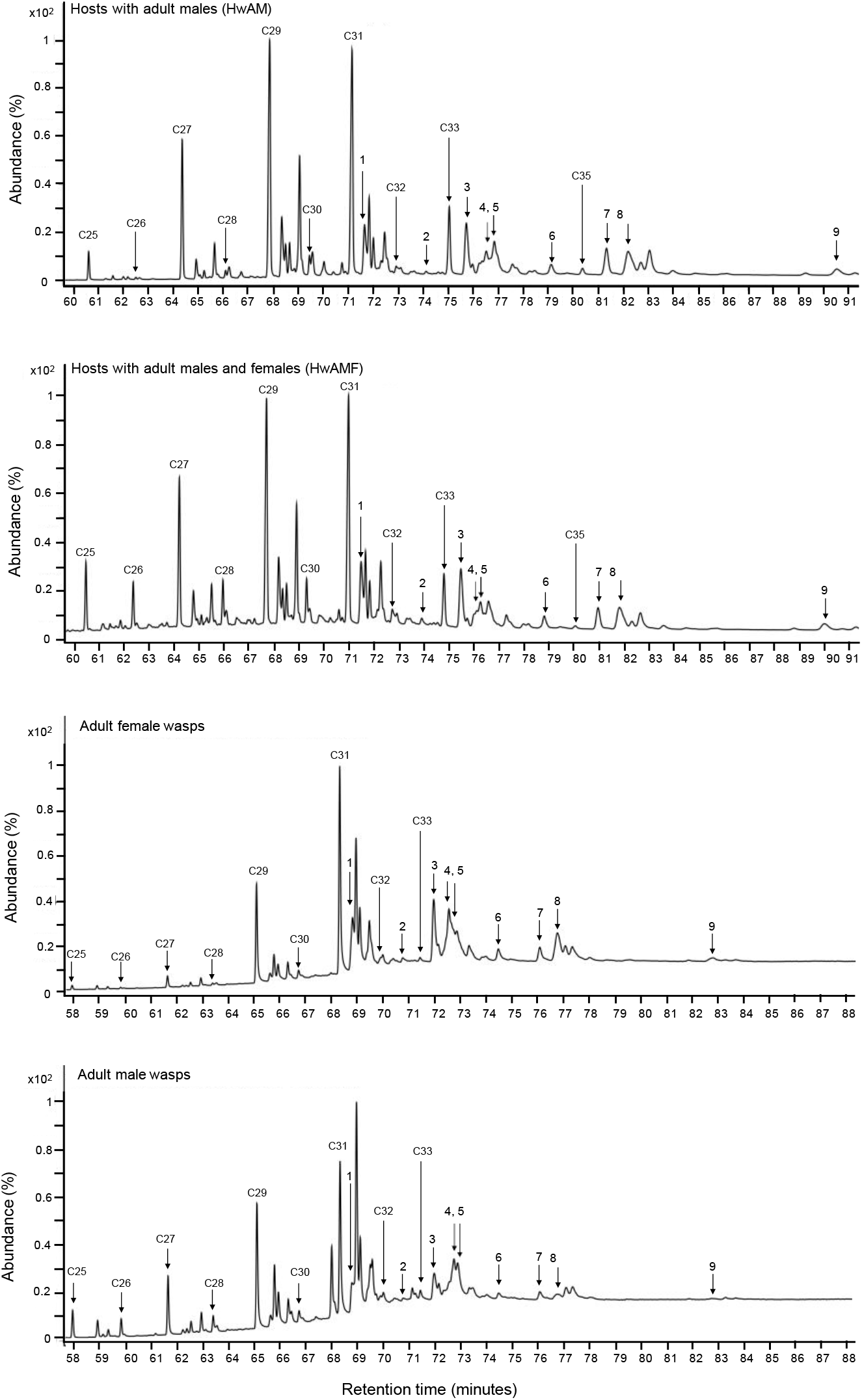
GC-MS profile obtained of different samples: Peak chromatogram of the non-polar fractions (enriched for CHCs) from HwAMF and HwAM shown in reference to the CHC profiles of adult male and female wasps. All the four samples share the same 47 CHC compounds (see also S.I., Table - S1). Straight chain alkanes (*n*C25- *n*C35) and nine compounds present in higher abundance in HwAMF and adult females (see also Table 1), are labelled (1-9) to their corresponding peaks.

**Figure 6:**
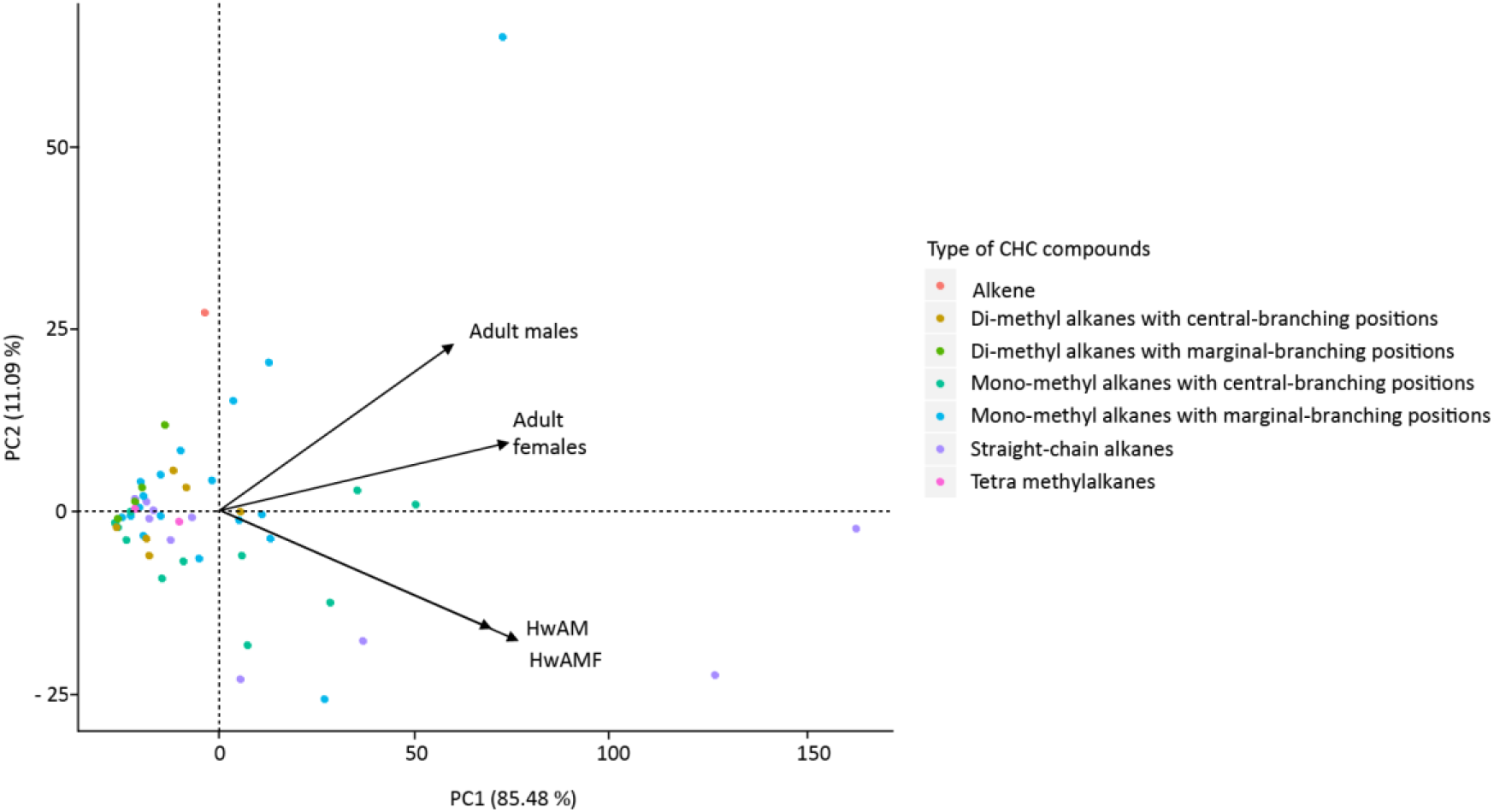
Principal component analysis of the cuticular hydrocarbon (CHC) profiles of different samples: A two-dimensional biplot of the principal component 1 and 2 explains 96.57% of the variance in the data. The samples HwAMF and HwAM show no separation unlike the adult males and females’ profiles.

The ability of males to distinguish HwAMF from HwAM (figure 2 b) could be due to some unique CHCs emanating from the former. Interestingly, the HwAMF and HwAM profiles share 47 compounds with both the adult males and female profiles. However, all the compounds differ in their respective relative abundances between HwAMF and HwAM as well as between adult males and females (Figure 8; SI - Table S1). Hence, it is likely that the males utilize the differences in relative abundances of compounds found in HwAMF as recognition signature CHCs.

**Figure 8:**
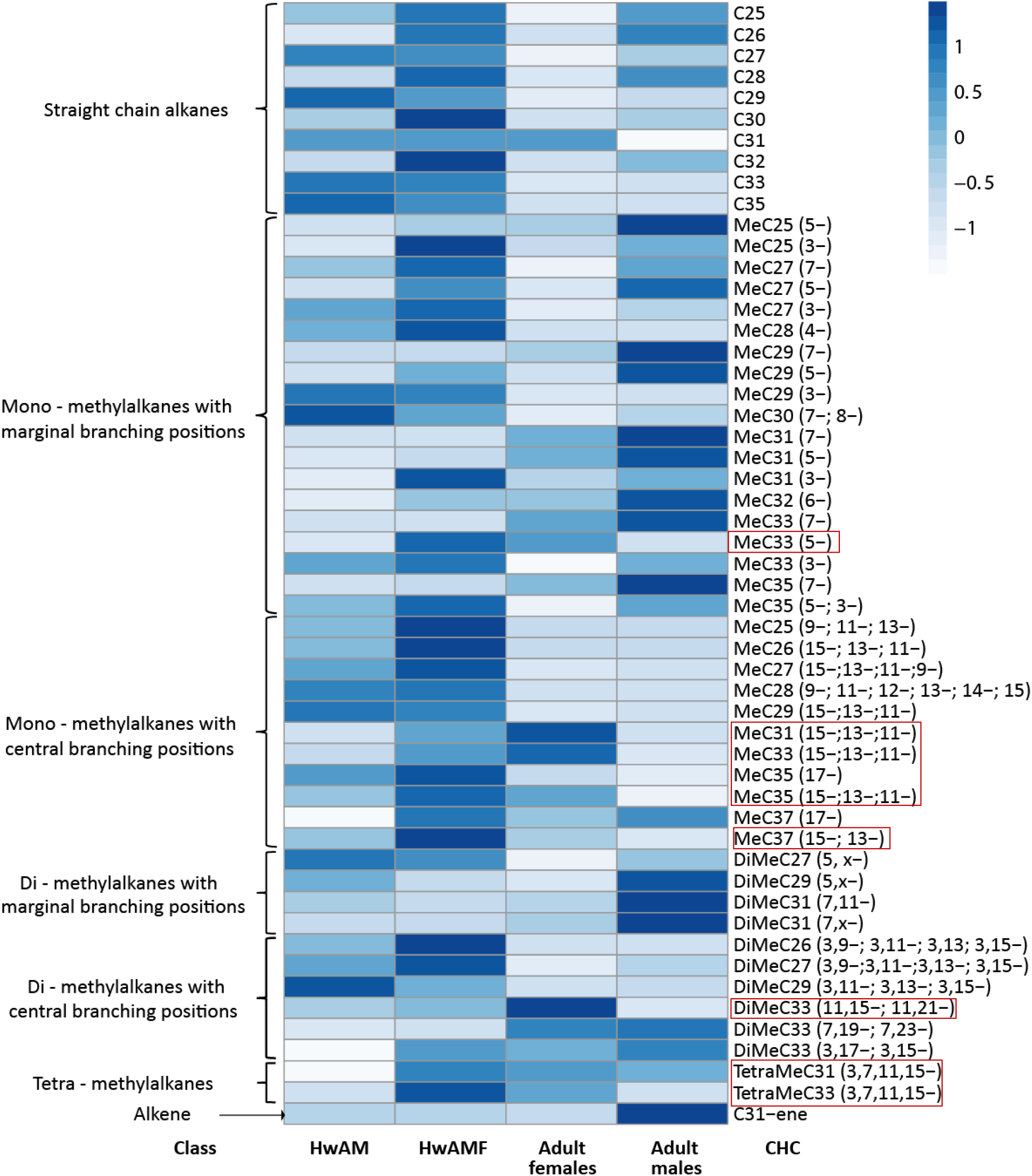
Heat map of CHCs: The heat map shows the abundance of various CHC compounds (scaled to the color intensity) in different samples. Names of the compounds are given on the right, the class they belong to is given on the left and sample names are given below. Compounds inside the red boxes have higher relative abundances (positive Cohen’s d, from Table 1) in HwAMF.

To investigate which signature CHCs are utilized by the males for detecting HwAMF, which contains both adult males and females, we tested whether males have the capability to distinguish between the adult male and female CHCs (see Methods). Males were given a choice between unparasitized hosts poured with CHC extract from adult females against those poured with the CHC extract from adult males. As figure 7 (a) shows, males prefer hosts with the CHC extract from adult females (*p* < 0.01, *r* = 0.6). This is not surprising as *Nasonia* males are known to utilize female CHCs for mate-recognition (Mair and Ruther, 2019; Buellesbach *et al*., 2018; Steiner *et al*., 2006). However, when their preference was checked for the adult male CHCs alone, males could not distinguish them from the solvent control (*p* = 0.2, *r* = 0.2; figure 7b) indicating that they do not utilize the male CHCs to detect hosts with adult females inside.

**Figure 7:**
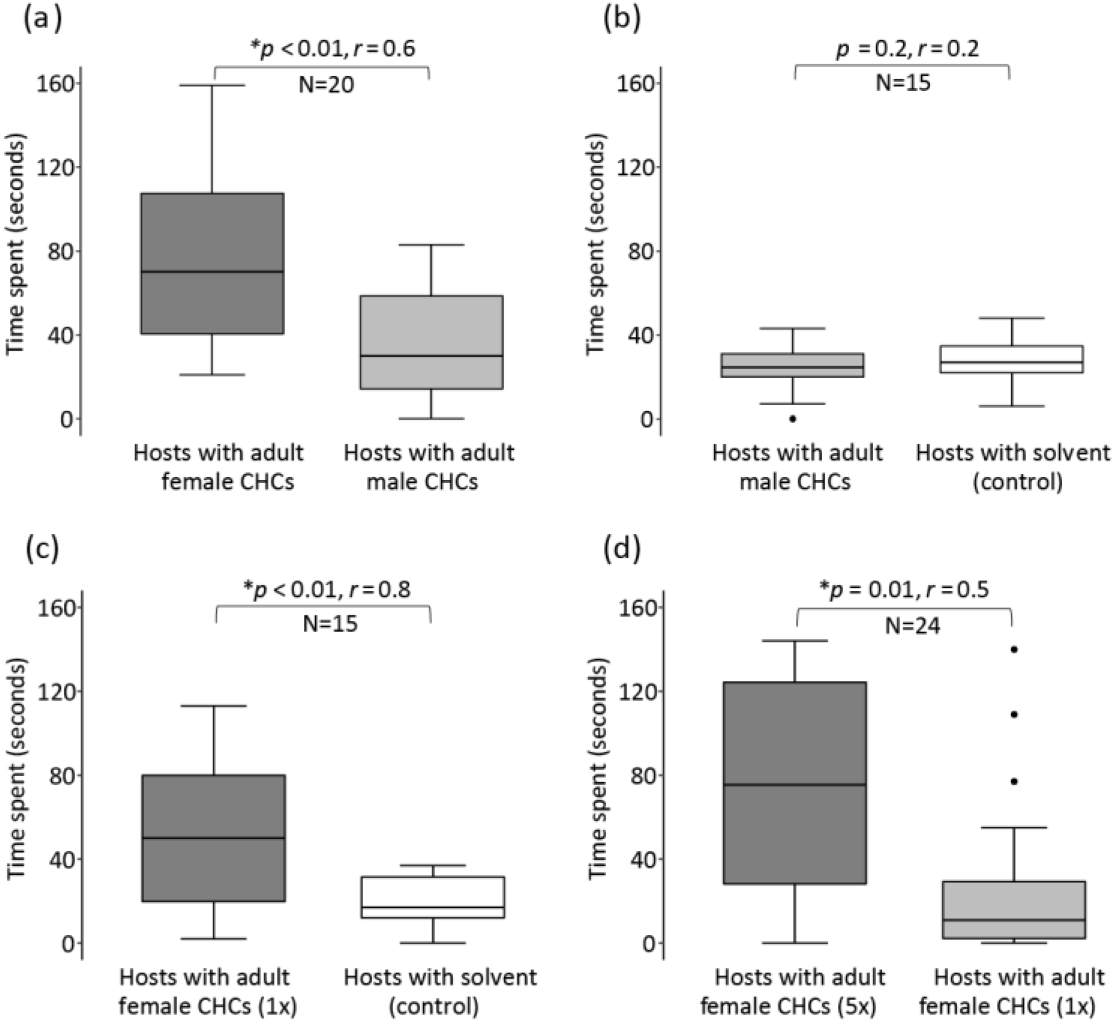
*N. vitripennis* males utilize cuticular hydrocarbons (CHCs) of females as olfactory cues to detect adult females within hosts: **(a)** Average time spent by males on the hosts poured over with female CHCs versus those with male CHCs. Males prefer the hosts poured over with adult female CHCs. **(b)** Average time spent by males on the hosts poured over with male CHCs versus the control poured with the solvent (Hexane). Males do not distinguish between the two types of hosts. Thus, males utilize female CHCs for detecting hosts with adult females inside. **(c)** Average time spent by males on the hosts poured over with female CHCs at 1x concentration versus control (Hexane). Males prefer the hosts poured over with female CHCs. **(d)** Average time spent by males on the hosts poured over with female CHCs at 1x concentration versus 5x concentration. Males prefer the hosts poured over with a higher concentration (5x) of adult female CHCs. The numbers above the boxes represent the *p-*value and the sample size (N), respectively. In boxplots, the horizontal bold line represents the median, boxes represent 25% and 75% quartiles and whiskers denote 1.5 interquartile ranges and black dots depict outliers. Statistical significance levels shown are according to Wilcoxon signed-rank test (statistically significant at *p* < 0.05) with (*) denoting a significant p-value. Wilcoxon effect size (*r*) values range from *r* = 0.1 - < 0.3 (small effect), *r* = 0.3 - < 0.5 (moderate effect) and *r* >= 0.5 (large effect).

As figure 2 (b) shows, males prefer the hosts with females (HwAMF) over those with all-male broods (HwAM). The preference for HwAMF can be easily explained by the presence of females inside and hence, female CHCs in HwAMF, possibly in higher concentration than in HwAM. It is not known, however, whether *N. vitripennis* males are capable of distinguishing different concentrations of CHCs as olfactory cues. To test this ability the males were given a choice between two different concentrations of the same female CHC extract. Three unparasitized hosts were poured over with 1x concentration of CHCs extract while the other three were poured over with 5x concentration. As figure 7 (d) indicates, males prefer hosts with a higher concentration of female-signature CHCs (*p* = 0.01, *r* = 0.5). This capability was further confirmed by showing male preference towards 1x concentration of female CHCs versus control (*p* < 0.01, *r* = 0.8; figure 7c). Therefore, the males have the ability to detect differences in concentration of the individual CHC compounds and identify hosts containing eclosed, but un-emerged, females.

### Which CHC component do the *N. vitripennis* males utilize to detect adult females within hosts?

It is plausible that males are utlilzing the relative abundance of various CHC compounds within the female profile for detecting HwAMF. Out of the 47 shared CHCs, only 9 compounds (Table 1; figure 8) have a higher (positive Cohen’s *d*) relative abundance in HwAMF (compared to HwAM) as well as the adult females (compared to males). These belong to different types of Mono-, Di-, and Tetra-methyl alkanes with chain length > *n*C30. It is likely that the higher relative abundances of these 9 compounds act as the olfactory cue for detecting HwAMF. This is consistent with the basic biology of *Nasonia* which exhibits a female-biased sex ratio indicating that female CHCs should have a higher abundance than male CHCs.

**Table 1:** List of candidate CHCs: Nine CHC compounds were found to have a higher (Cohen’s *d*) relative abundance in the adult females than in males, similar to that found in HwAMF over HwAM. The major class of compounds are the mono-methyl alkanes (with central-branching positions), Di- and Tetra-methyl alkanes with the carbon chain length > 30. (For a full list of identified CHCs, see S.I. - Table S1).

| S.No. | Compound name | Class | Effect size (Cohen's *d*) |  |
| --- | --- | --- | --- | --- |
|  |  |  | Between HwAMF and HwAM | Between Adult females and males |
| 1 | MeC33 (5-) | Monomethylalkanes (with central-branching positions) >C30 | 1.83 | 1.91 |
| 2 | MeC31 (15-;13-;11-) |  | 3.98 | 6.52 |
| 3 | MeC33 (15-;13-;11-) |  | 2.60 | 2.52 |
| 4 | MeC35 (17-) |  | 0.95 | 0.49 |
| 5 | MeC35 (15-;13-;11-) |  | 1.87 | 2.81 |
| 6 | MeC37 (15-;13-) |  | 1.77 | 3.39 |
| 7 | DiMeC33 (11,15-;11,21-) | Dimethylalkane (with central-branching positions) >C30 | 2.20 | 3.62 |
| 8 | TetraMeC31 (3,7,11,15-) | Tetramethylalkanes | 7.90 | 0.31 |
| 9 | TetraMeC33 (3,7,11,15-) |  | 3.05 | 1.11 |

## Discussion

Our study shows that *N vitripennis* males can seek out adult females inside the fly host using olfactory cues emanating from the females still inside the puparium. That males are attracted by female CHCs is not surprising. But what is remarkable is the males’ ability to utilize the olfactory signature within the female CHC profile to detect hosts about to release adult females. Thus, it shows the presence of a solid, and as yet unknown, mate-finding strategy of *N. vitripennis* males. More surprisingly, this ability is restricted to *N. vitripennis* despite a very similar habitat and ecology for all the four *Nasonia* species. One possibility of why the other species, especially *N. giraulti*, do not show this ability is the phenomenon of within-host mating, where mating happens within the fly host before emergence. However, this still does not explain why the *N. longicornis* and *N. oneida* males do not have this ability as they show intermediate rates of within-host-mating (Giesbers *et al*., 2016).

Males of all the four *Nasonia* species share the ability to distinguish parasitized hosts from unparasitized ones with other parasitoid wasps like *Pimpla disparis* (Hrabar *et al*., 2012; Danci *et al*., 2014), *Lariophagus distinguendus* (Steiner *et al*., 2005) and *Cephalonomia tarsalis* (Collatz *et al*., 2009). But whether this ability also extends to distinguishing hosts containing females from those that do not, like *N. vitripennis*, is not clear. Therefore, the present study is one of the first reports of this mate-finding strategy employed by *N. vitripennis* males.

The capability of *N. vitripennis* males to distinguish between different concentrations of female CHCs (figure 7 d) underscore their ability to find, even in a patch, hosts with varying number of females inside. Assuming that the cues increase in intensity with the number of females inside a host, a male can now seek out hosts with the maximum number of females. This capability has the potential to further increase individual male fitness. Moreover, this ability can also explain why males can distinguish between HwL from unparasitized ones (Figure 1 a). In the former case, the males are probably detecting the olfactory cues left behind by the parasitizing females. But these cues get swamped out when given a choice with HwAMF as it usually contains several females inside. This ability can also potentially bring several males in contact with each other resulting in increased male-male conflict and then trigger selection for more aggressive male behaviour, both for access to females and territoriality. There is some evidence that this could have happened as *N. vitripennis* males are the most aggressive among the four species (Leonard and Boake, 2006; Giesbers *et al*., 2016; Mair and Ruther, 2018).

Another curious phenomenon found in this study is the compositional uniformity of CHCs from hosts with and without females (Figure 8; SI - Table S1). This finding is consistent with other studies reporting such uniformity even in adult males and females (Ruther *et al*., 2011; Steiner *et al*., 2005). Yet, a male *N. vitripennis* can still detect adult females within hosts. Therefore, the most parsimonious explanation for this behaviour is the ability of males to detect variations of the individual CHC components from the two types of hosts and use that as female specific signature cue. We have analysed these variations across the adult male and female CHCs and have hypothesized a specific list serving as female specific signature (Table 1) which awaits further empirical validation.

The *Nasonia* genus represents one of the best-characterized insect model systems for understanding the chemical and behavioral basis of communication between the sexes (Mair *et al*., 2019). Despite this accrued information, our study uncovers a previously unknown male mate-finding strategy in *N. vitripennis*. Moreover, *Nasonia* belongs to the superfamily Chalcidoidea which has an estimated 500,000 species (Heraty *et al*., 2013), making it one of the most speciose of any animal group. Many of these species share a similar idiobiont lifestyle with *Nasonia*. Even if a fraction of these species share the ability to detect females still inside their hosts, then this mate-finding strategy can be one of the most widespread in the animal kingdom.

## Materials

### Fly host used

All *Nasonia* cultures were raised on pupae of the fly, *Sarcophaga dux*, which has a life-cycle of 11 days at 25°C. The fly larvae were fed with chicken liver, and the pupae were stored at 4°C. The fly pupae kept at 4°C for ≤48 hours were used in all the experiments and are designated as ‘unparasitized’ hosts.

### *Nasonia* strains used

The wasp strains of the four *Nasonia* species used were NV-IPU08 (a field strain obtained from Punjab, India) for *N. vitripennis*, NL-MN8501 for *N. longicornis*, NG-RV2XU for *N. giraulti*, and NO-NY11/36 for *N. oneida*. These were reared in a 24-hour light cycle at 25°C and 60% relative humidity and had an average life-cycle of 14 days for *N. vitripennis*, 14.5 days for *N. longicornis*, 15 days for *N. giraulti*, and 16 days for *N. oneida*. The different life-stages include egg (1-2 days), larva (2-7 days), pupa (8-12 days), and adult (13-16 days) (Whiting, 1967). One female (either mated or virgin) was provided with two unparasitized hosts for 48 hours and then removed. The parasitized hosts were either kept for wasps’ emergence or used in the experiments as required. All experiments were done using parasitized hosts containing larval wasps (two-day post-parasitization) or eclosed adult wasps inside (thirteen-/fourteen/fifteen-days, depending on the species). The former has been abbreviated as HwL (hosts with larval wasps) and the latter as HwAMF (hosts with adult males and females)/HwAM (hosts with adult all-male broods). Experiments to investigate which cues are utilized by males were done with *N. vitripennis* (NV-IPU08).

### Behavioural assay and determination of cues used

To test which type of host a male preferred, a cafeteria arena having two concentric circles (outer 9 cm and inner 5 cm diameter) divided into six equal chambers was printed on a white sheet of paper over which a glass Petri plate (sterilized with ethanol, then with HPLC grade n-hexane and autoclaved) was placed. Autoclaved distilled water was added along the circumference to prevent males from escaping. This setup was placed on a wooden platform with a 5-watt LED lamp placed 30 cm above it. Each new male assay, *i.e*., every data point, was obtained using a fresh set of six hosts and a fresh Petri plate. Each data point was obtained by randomly choosing a single virgin male (<48 hours old) from all-male broods to prevent any sensory bias accumulating because of co-development with females. Each by a video camera (Logitech C615 HD webcam) at 25°C ±1°C. Each male was used only once and then discarded to prevent prior experience influencing their preference.

All parasitized hosts were handled with separate sets of forceps (sterilized with 70% ethanol, HPLC grade *n*-hexane, and autoclaved). Male preference for either type of hosts (SI – Figure S1) was quantified by the average time spent on each host for the first 4 minutes. The time spent was counted from when a male climbed onto a host and continued till it dismounted and abandoned it. All parasitized hosts were cracked open after the experiment to check whether all had the requisite sex, developmental stage as well as alive or dead wasps. The presence of emerged adult wasps inside was insured by using hosts just one day before emergence, *i.e*., 13 days for *N. vitripennis* and *N. longicornis*, 14 days for *N. giraulti* and 15 days for *N. oneida*. Care was taken to note the absence of any emergence holes made by the adult wasps within the hosts. If not, then the entire data point was discarded from further analysis. A control experiment to check for the males’ inherent directional bias was done using all six unparasitized hosts—none of the four species showing any such directional bias (SI - Figure S2).

- **Auditory cues:** To investigate any possible auditory cues coming from the adult wasps, HwAMF were freeze-killed by keeping them at −80°C for 2 hours, and then brought at room temperature, which was confirmed by an LCD digital I.R. temperature laser gun (Dual Laser Optical Focus Temperature Gun, NUB8580) and used in the experiment within 2 hours.
- **Visual cues:** To check for progressive darkening of the puparial halves serving as a visual cue, the anterior half of the puparia of the unparasitized hosts of different ages and different degrees of darkening were used (SI - Figure S3).
- **Olfactory cues:** To check whether olfactory cues are used, male preference was recorded towards puparial halves from the anterior part of the parasitized hosts of varying ages (HwAMF and HwL) against those of unparasitized ones of the same age. Male preference was also tested for the total extract of the puparial halves obtained through Dichloromethane (DCM) extraction and the non-polar and polar fractions enriched through column chromatography (see below). Extracts enriched for cuticular hydrocarbons (CHCs) from both adult male and female wasps were obtained using the 50 individuals of each, processed through column chromatography, and then used to test the behavioural response of the males.

### Column Chromatography Method

#### a) Chemical extraction of puparial halves

Puparial halves (n=50) obtained from HwAMF were extracted using 1 ml of HPLC grade *n*-hexane (Merck Corp.) in a glass vial at room temperature. This extract was poured into a column made of glass Pasteur pipettes (inner diameter = 0.7 cm) packed with baked glass wool and 3 cm of activated silica gel (100-200 Mesh; Merck Millipore). The non-polar compounds were eluted in *n*-hexane (3/8 dead volume), followed by the polar compounds’ elution with a Dichloromethane and Methanol solution (9:1). Both the polar and non-polar fractions were concentrated to 50 μl with a Nitrogen stream. Puparial halves obtained from HwAM were extracted through the same protocol and fractionated to a non-polar fraction.

#### b) Extraction of CHCs from the adult wasps

50 individuals were dipped separately in two glass vials with 500 μl of HPLC grade n-hexane (Merck Corp.) 10 minutes. The extract was pipetted out into a fresh set of glass vials. The extract was poured into a column made out of glass Pasteur pipettes (inner diameter = 0.7 cm) packed with baked glass wool and 1.5 cm of activated silica gel. The non-polar fraction enriched in CHCs was eluted in *n*-hexane (3/8 dead volume) and concentrated to 250 μl under a Nitrogen stream for both males and females separately.

Another set of extraction of adult female CHCs was done through the same protocol and concentrated to 50 μl for use as 5X concentrated fraction of CHCs (Figure 7 d).

### Gas Chromatography-Mass Spectrometry (GC-MS)

For identification of the chemicals, the non-polar fraction of the extract obtained from the puparial halves of HwAM, HwAMF (2 μl of each) as well as the extract from 2 individuals each of both adult males and females (separately dipped in 20 μl of Hexane for 10 minutes and concentrated to 2 μl under Nitrogen stream), were all separately injected (split-less mode) into a gas-chromatograph coupled with Mass spectrometer (Agilent 7890B, 5977C GC-MS). The machine had a capillary column, HP-5MS (Agilent J&W), with an operational mode of electron impact ionization at 70eV (Quadrupole temperature of 150°C). The inlet temperature and the auxiliary line temperature were maintained at 320°C, and Helium was used as the carrier gas with an avg. velocity = 36.2 cm/sec. The oven temperature was programmed from 40°C with a hold of 5 minutes, increased from 40°C to 300°C at 4 °C/min with a final hold for 25 minutes.

CHC compounds were identified according to their characteristic diagnostic ions and resulting mass spectra (Lockey, Kenneth H., 1988; Howard, Ralph W., 1993; Ruther, J. et al., 2011; Carlson, D. A. *et al*., 1999). The branched-chain alkanes, resulting from mass fragmentations at branching points, were identified with the extracted ion chromatogram (EIC-*m/z*) and by comparing the retention index values with the literature data (Steiner, S. *et al*., 2006; Buellesbach, J. *et al*., 2018). An *n*-alkane (C8-C40, SUPELCO) standard was also analyzed under the same conditions to calculate the relative retention indices to characterize the CHCs (Van Den Dool, H. and P. Dec Kratz, 1963; Carlson, D. A. *et al*., 1998). Peaks were analyzed in Mass Hunter Workstation Software vB.08.00 (Agilent Technologies). For calculating the relative abundance of each identified peak, each was divided by the area of the most abundant peak within each sample (*i.e*., *n*C-29 in HwAM, *n*C-31 in HwAMF, as well as adult female and MeC31 (7-) in adult males). The peak ratios relative to the highest peak (taken as 100 %) were transformed into percentages for subsequent statistical analysis.

### Statistical analysis

All statistical analysis was done in RStudio, v1.2.5033 (RStudio Team, 2015). Shapiro-Wilk test (Shapiro, Samuel Sanford, and Martin B. Wilk., 1965) was used to test for normality using the *stats* package (R Core Team, 2020). The obtained data tested negative for normality; hence, Wilcoxon signed-rank test (significant at *p* < 0.05) was used to assess male preference in all the assays. Wilcoxon effect size (*r*) was calculated from the *Z*-statistic obtained from the Wilcoxon signed-rank test using the *stats* package. Boxplots were made by using the *ggplot2* package (H. Wickham., 2016). Heatmap was made using the *pheatmap* (Raivo Kolde., 2019) package in R. Principal Component Analysis was done using the *ggplot* and *ggfortify* (Horikoshi M. and Li W., 2016; Horikoshi M. and Tang Y., 2018) package in R.

## Acknowledgments

We thank Ruchira Sen and all Evogen lab members for comments on the study and Leo Beukeboom (University of Groningen) for providing us with strains of the four *Nasonia* species.

## Author contributions

GP conceived the study, designed the experiments, collected data and analysed the results. AY and GP performed the GC-MS. AA supervised the GC-MS. AS helped with data collection. GP and RR wrote the manuscript.

## Data availability

All behavioral assays are available as videos on https://www.youtube.com/channel/UCBh3wyHrAty7dvcLNX6HeOw/videos.

