## Supplementary Information for "The males of the parasitoid wasp, *Nasonia vitripennis*, can identify which fly hosts contain females"

**This file includes:**

Figures S1 to S3
Table S1
Legends for figures S1 to S3
Legend for Table S1

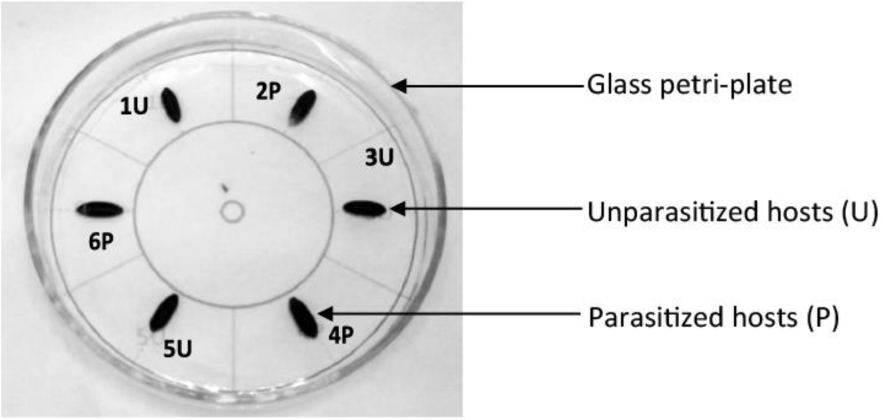


**Figure S1. Cafeteria arena used for all the behavioural assays:** The arena had two concentric circles (the outer 9 cm and the inner 5 cm diameter, respectively) divided into six equal chambers printed on a white sheet of paper over which a glass Petri plate (sterilized with ethanol, then with HPLC grade n-hexane and autoclaved) was placed. Autoclaved distilled water was added along the circumference to prevent males from escaping. This setup was placed on a wooden platform with a 5-watt LED lamp placed 30 cm above it. Each male was dropped at the centre of this arena, and the behaviour was recorded for 4 minutes by a video camera (Logitech C615 HD webcam) at 25°C ±1°C.

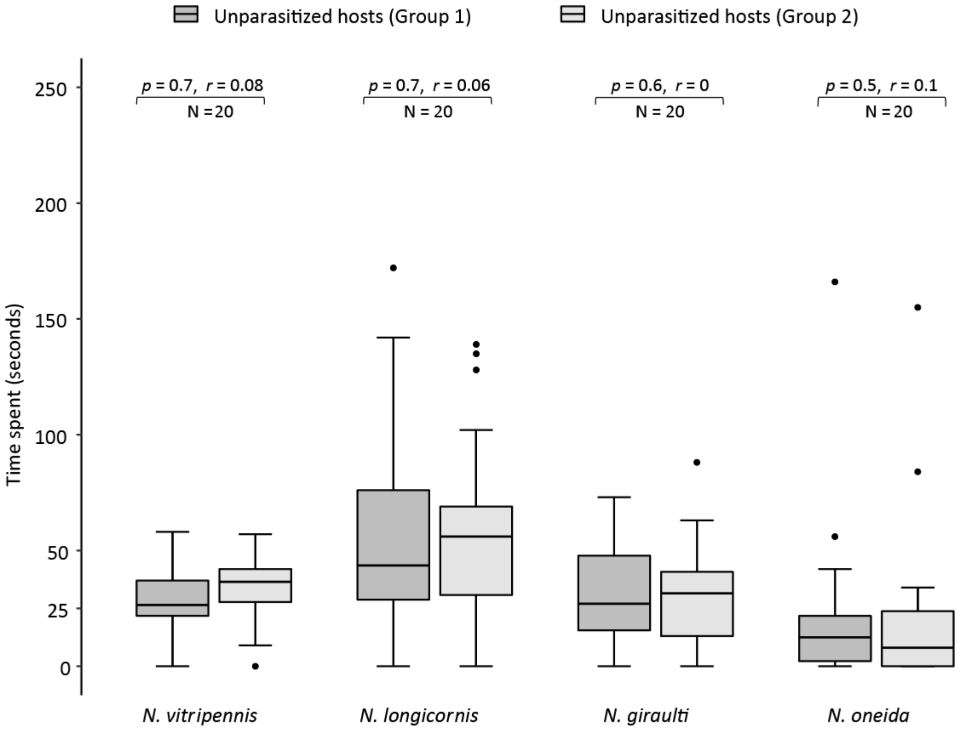


**Figure S2. No inherent directional bias of male *Nasonia* in the cafeteria arena:** To check whether *Nasonia* have an inherent bias towards any one direction within the arena, males were given a choice between six hosts (all unparasitized) and scored for time spent on each. These six unparasitized hosts were randomly divided into 2 groups of three each, to mimic the actual choice assay (as in figure S1), with the help of random numbers assigned to each (from 0 to 1). Random numbers with the lowest three values for each assay (N = 20) were assigned to "group 1" and the remaining to "group 2". Wilcoxon- signed-rank test was performed for assessing the difference in distribution of the two groups of unparasitized hosts. As the figure above indicates, no significant difference was found for the two groups across all the four species, indicating no evidence of a directional bias.

The numbers above the boxes represent the *p*-value and the sample size (N) for each species. In boxplots, the horizontal bold line within each box represents the median, boxes represent 25% and 75% quartiles, whiskers denote 1.5 interquartile ranges and black dots depict outliers. Statistical significance levels shown are according to Wilcoxon signed-rank test (statistically significant at $p < 0.05$) with (∗) denoting a significant *p*-value. Wilcoxon effect size (*r*) values range from $r = 0.1 - < 0.3$ (small effect), $r = 0.3 - < 0.5$ (moderate effect) and $r >= 0.5$ (large effect). Species names are given at the bottom.

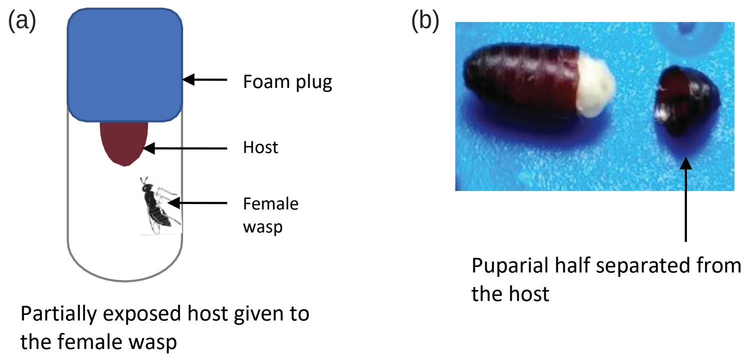


**Figure S3. Use of puparial halves for the assays:** A) Individual unparasitized host was half embedded in foam plugs put in vials so that only a part (anterior part) of the host was accessible to the female for parasitization. Half of such vials were exposed to individual mated female wasps while others remained female free, hence, unparasitized. B) After 48 hours, the hosts were taken out from plugs, carefully cracked open from the anterior part and the puparium separated from it. The hosts that were exposed to mated females were checked for the presence of eggs (now localized at the head of the fly pupa), and those found without eggs were discarded. The puparia of HwAMF and hosts containing adult fly (ten-day old) were collected after confirming the presence of adult wasps and adult fly, respectively.

**Table - S1: List of all identified CHC compounds:** There are 53 identified CHCs in HwAMF and HwAM and 47 in adult males and females (N = 4 for each sample). Each value represents mean relative abundance (± SEM or standard error of mean) expressed as a percentage. Retention times mentioned are of the samples HwAMF and HwAM (for chromatograms see figure 5).

| S.No. | Retention Time | Linear Retention Index | Compound name | Mean relative abundance (%) ± SEM | | | |
|---|---|---|---|---|---|---|---|
| | | | | HwAM | HwAMF | Adult female | Adult male |
| 1 | 60.6 | 2500 | C25 | 9.40 ± 0.58 | 14.88 ± 2.27 | 4.35 ± 0.35 | 12.72 ± 0.25 |
| 2 | 61.1 | 2534 | MeC25 (9-; 11-; 13-) | 0.73 ± 0.19 | 2.57 ± 0.15 | - | - |
| 3 | 61.6 | 3041 | MeC25 (5-) | 1.72 ± 0.30 | 3.04 ± 0.41 | 2.96 ± 0.26 | 8.08 ± 0.26 |
| 4 | 62 | 3141 | MeC25 (3-) | 1.29 ± 0.26 | 4.65 ± 0.29 | 1.70 ± 0.17 | 2.89 ± 0.09 |
| 5 | 62.5 | 2600 | C26 | 0.87 ± 0.17 | 8.76 ± 2.07 | 0.91 ± 0.08 | 7.89 ± 0.12 |
| 6 | 62.66 | 3355 | DiMeC26 (3,9-; 3,11-; 3,13; 3,15-) | 0.87 ± 0.18 | 2.35 ± 0.05 | - | - |
| 7 | 63.15 | 2635 | MeC26 (15-; 13-; 11-) | 0.54 ± 0.09 | 1.70 ± 0.17 | - | - |
| 8 | 64.4 | 2700 | C27 | 46.42 ± 0.51 | 43.62 ± 3.14 | 8.83 ± 0.59 | 27.30 ± 0.63 |
| 9 | 64.9 | 2735 | MeC27 (15-;13-;11-;9-) | 8.61 ± 0.73 | 14.05 ± 0.21 | 0.82 ± 0.06 | 1.69 ± 0.16 |
| 10 | 65.07 | 3149 | MeC27 (7-) | 1.36 ± 0.11 | 2.32 ± 0.10 | 0.67 ± 0.08 | 1.74 ± 0.05 |
| 11 | 65.25 | 3173 | MeC27 (5-) | 2.82 ± 0.29 | 4.48 ± 0.33 | 2.59 ± 0.16 | 5.04 ± 0.06 |
| 12 | 65.66 | 3244 | MeC27 (3-) | 13.15 ± 0.70 | 17.55 ± 0.94 | 4.84 ± 0.37 | 8.75 ± 0.20 |
| 13 | 65.8 | 2785 | DiMeC27 (5, x-) | 1.28 ± 0.06 | 1.21 ± 0.11 | 0.54 ± 0.10 | 0.93 ± 0.09 |
| 14 | 66.1 | 2800 | C28 | 2.76 ± 0.35 | 9.74 ± 1.51 | 1.50 ± 0.07 | 7.97 ± 0.30 |
| 15 | 66.25 | 3365 | DiMeC27 (3,9-;3,11-;3,13-; 3,15-) | 5.47 ± 0.50 | 7.92 ± 0.26 | 1.52 ± 0.12 | 3.38 ± 0.27 |
| 16 | 66.67 | 2832 | MeC28 (9-; 11-; 12-; 13-; 14-; 15) | 3.68 ± 0.46 | 3.88 ± 0.39 | - | - |
| 17 | 67.1 | 3340 | MeC28 (4-) | 0.99 ± 0.16 | 2.08 ± 0.15 | - | - |
| 18 | 67.85 | 2900 | C29 | 100.00 ± 0 | 88.01 ± 1.11 | 53.79 ± 1.71 | 63.47 ± 1.26 |
| 19 | 68.35 | 2932 | MeC29 (15-;13-;11-) | 27.88 ± 1.21 | 27.28 ± 0.49 | 5.31 ± 0.21 | 7.06 ± 0.33 |
| 20 | 68.5 | 3349 | MeC29 (7-) | 10.62 ± 0.69 | 10.94 ± 0.23 | 12.81 ± 0.64 | 29.72 ± 0.03 |
| 21 | 68.65 | 3373 | MeC29 (5-) | 10.56 ± 0.62 | 13.74 ± 0.29 | 10.64 ± 2.47 | 17.32 ± 0.36 |
| 22 | 69.05 | 3537 | MeC29 (3-) | 43.45 ± 0.98 | 40.60 ± 0.52 | 8.75 ± 0.38 | 13.12 ± 0.42 |
| 23 | 69.17 | 2987 | DiMeC29 (5,x-) | 3.44 ± 0.08 | 2.09 ± 0.12 | 1.48 ± 0.12 | 5.53 ± 0.20 |
| 24 | 69.45 | 3000 | C30 | 6.56 ± 0.52 | 13.06 ± 0.72 | 5.00 ± 0.16 | 6.44 ± 0.52 |
| 25 | 69.58 | 3395 | DiMeC29 (3,11-; 3,13-; 3,15-) | 10.26 ± 0.30 | 5.74 ± 0.31 | 1.24 ± 0.03 | 1.98 ± 0.20 |
| 26 | 70 | 3560 | MeC30 (7-; 8- ) | 6.96 ± 0.53 | 4.97 ± 0.26 | 1.73 ± 0.19 | 2.88 ± 0.59 |

| S.No. | Retention Time | Linear Retention Index | Compound name | Mean relative abundance (%) ± SEM | | | |
|---|---|---|---|---|---|---|---|
| | | | | HwAM | HwAMF | Adult female | Adult male |
| 27 | 70.77 | 3453 | C31-ene | 4.19 ± 0.45 | 3.49 ± 0.23 | 1.43 ± 0.20 | 41.43 ± 2.54 |
| 28 | 71.15 | 3100 | C31 | 98.98 ± 0.46 | 100.00 ± 0 | 100.00 ± 0 | 81.25 ± 1.11 |
| 29 | 71.65 | 3131 | MeC31 (15-;13;11-) | 26.90 ± 0.99 | 33.24 ± 0.54 | 39.54 ± 0.75 | 26.82 ± 1.15 |
| 30 | 71.85 | 2551 | MeC31 (7-) | 26.70 ± 0.89 | 25.37 ± 0.83 | 55.15 ± 1.02 | 100.00 ± 0 |
| 31 | 72 | 2573 | MeC31 (5-) | 11.13 ± 0.64 | 13.86 ± 0.38 | 23.09 ± 0.84 | 35.04 ± 0.46 |
| 32 | 72.3 | 3171 | DiMeC31 (7,11-) | 3.31 ± 0.43 | 2.60 ± 0.33 | 3.14 ± 0.13 | 7.66 ± 0.40 |
| 33 | 72.45 | 2740 | MeC31 (3-) | 14.60 ± 0.51 | 26.37 ± 1.28 | 17.61 ± 0.39 | 20.71 ± 1.36 |
| 34 | 72.57 | 3178 | DiMeC31 (7,x-) | 3.07 ± 0.21 | 2.84 ± 0.16 | 4.83 ± 0.39 | 18.54 ± 1.04 |
| 35 | 72.9 | 3200 | C32 | 3.67 ± 0.30 | 7.44 ± 0.41 | 3.58 ± 0.13 | 4.92 ± 0.45 |
| 36 | 73.2 | 2750 | MeC32 (6-) | 2.62 ± 0.26 | 4.16 ± 0.29 | 4.20 ± 0.12 | 6.45 ± 0.17 |
| 37 | 74.1 | 3074 | TetraMeC31 (3,7,11,15-) | 1.28 ± 0.05 | 4.20 ± 0.26 | 3.71 ± 0.67 | 3.28 ± 0.72 |
| 38 | 75.05 | 3300 | C33 | 29.64 ± 0.41 | 27.96 ± 1.58 | 2.60 ± 0.29 | 6.49 ± 0.09 |
| 39 | 75.7 | 3335 | MeC33 (15-;13-;11-) | 34.69 ± 0.55 | 42.78 ± 2.13 | 47.27 ± 3.08 | 31.86 ± 3.03 |
| 40 | 75.95 | 2775 | MeC33 (7-) | 4.38 ± 0.12 | 4.20 ± 0.21 | 7.50 ± 0.66 | 10.23 ± 0.15 |
| 41 | 76.22 | 2858 | MeC33 (5-) | 5.30 ± 0.22 | 7.78 ± 0.93 | 6.98 ± 0.35 | 5.47 ± 0.43 |
| 42 | 76.54 | 2810 | DiMeC33 (11,15-;11,21) | 14.23 ± 0.52 | 16.59 ± 0.56 | 25.45 ± 2.51 | 10.33 ± 1.55 |
| 43 | 76.7 | 3012 | DiMeC33 (7,19-; 7,23-) | 5.26 ± 0.18 | 5.71 ± 0.40 | 11.09 ±0.31 | 11.13 ±0.99 |
| 44 | 76.85 | 2939 | MeC33 (3-) | 22.41 ± 0.83 | 27.03 ± 1.13 | 10.05 ± 1.22 | 21.64 ± 1.12 |
| 45 | 77.6 | 2610 | DiMeC33 (3,17-; 3,15-) | 5.79 ± 0.14 | 11.30 ±1.40 | 10.50 ± 1.18 | 11.92 ±1.38 |
| 46 | 79.25 | 3260 | TetraMeC33 (3,7,11,15-) | 6.57 ± 0.19 | 12.06 ± 1.26 | 9.48 ± 1.48 | 6.74 ± 0.91 |
| 47 | 80.4 | 3500 | C35 | 3.38 ± 0.13 | 2.68 ± 0.34 | - | - |
| 48 | 81.3 | 3524 | MeC35 (17-) | 18.59 ± 0.80 | 22.02 ± 2.42 | 14.03 ± 2.16 | 12.06 ± 1.82 |
| 49 | 82.2 | 3532 | MeC35 (15-;13-;11-) | 26.47 ± 1.73 | 40.08 ± 4.83 | 31.74 ± 4.33 | 12.63 ± 2.08 |
| 50 | 82.7 | 2950 | MeC35 (7-) | 5.50 ± 0.99 | 6.22 ± 0.42 | 9.41 ± 1.15 | 15.80 ± 1.93 |
| 51 | 83.05 | 2971 | MeC35 (5-; 3-) | 16.31 ± 1.72 | 21.11 ± | 2.46 | 10.57 ± 1.89 |
| 52 | 89.3 | 3730 | MeC37 (17-) | 1.96 ± 0.05 | 3.01 ± 0.51 | 2.55 ± 0.45 | 2.85 ± 0.41 |
| 53 | 90.5 | 3722 | MeC37 (15-; 13-) | 8.65 ± 0.48 | 16.34 ± 3.04 | 7.88 ± 0.70 | 3.82 ± 0.48 |
